# A Sexually Dimorphic Neuronal Cluster in the Mouse Medial Amygdala Exhibits Binary Activation Mode Based on Male Sexual Status

**DOI:** 10.1101/2025.07.09.663831

**Authors:** Tamar Licht, Adan Akarieh, Aya Dhamshy, Dan Rokni

**Affiliations:** Department of Medical Neurobiology, Faculty of Medicine and IMRIC, The Hebrew University of Jerusalem, Jerusalem, Israel

## Abstract

Increasing scientific interest has been directed toward understanding sexual dimorphism in the brain. Although various brain structures exhibit masculine or feminine characteristics, no strictly binary anatomical feature, such as those seen in genitalia. has been identified. In this study, we identified a dense, sexually dimorphic cluster of neurons in the posterodorsal medial amygdala (MeApd), which we named DIMPLE, that exhibited a remarkable binary pattern of cFos activation. Using the TRAP2 (Targeted Recombination in Active Populations) transgenic mouse model, we found that it was consistently labeled in all females, regardless of age or sexual experience. In males, however, DIMPLE was not labeled in any of the adult virgins but was evident pre-weaning and following mating. Surgical removal of gonads (ovariectomy or orchiectomy) did not alter the labeling pattern of DIMPLE in either sex. Interestingly, a single intraperitoneal injection of prolactin, a hormone known to increase in males after mating, induced DIMPLE labeling in virgin males. However, treatment with cabergoline, a potent inhibitor of prolactin secretion, did not prevent DIMPLE labeling in females or in post-mating males. Given the established role of the MeApd in social and reproductive behaviors, we propose that DIMPLE may support neural mechanisms underlying female-typical behavior and potentially contribute to post-mating behavioral shifts in males.

Graphical abstract:
Summary of DIMPLE activity during puberty and sexual status.

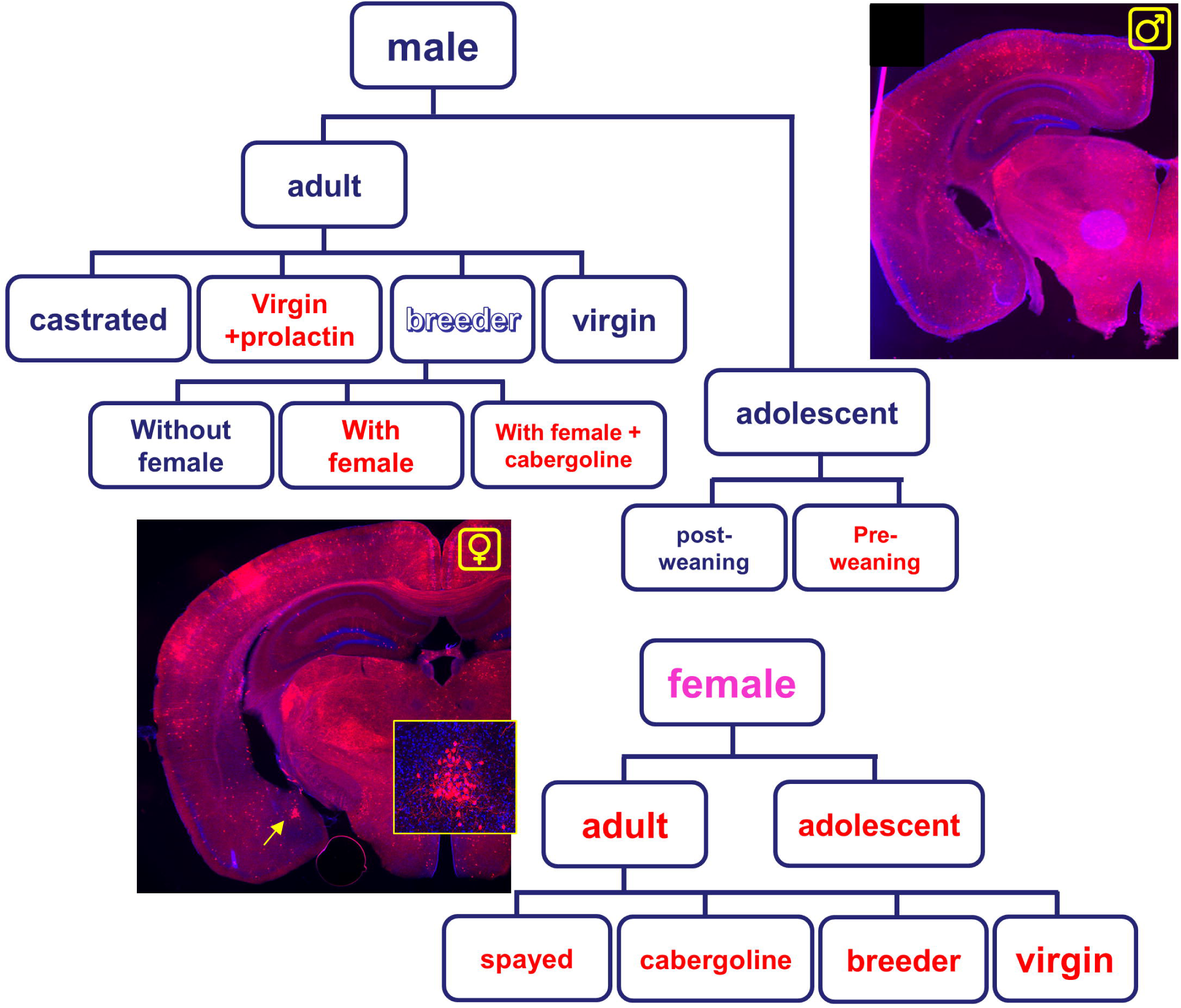

## Introduction

Genitalia, body shape, and hormones exhibit clear sex differences. Dimorphism in the brain, however, is still a controversial topic. The sex differences that have been found in the human brain so far, manifest as overlapping distributions rather than clear binary features, exclusive to one of the sexes^1–3^. Human and animal studies have demonstrated sex differences in anatomy, connectivity, neuronal activity, paracrine and endocrine signaling, and gene expression^4–11^, but these are mostly gradual and overlapping. Clear binary differences in the brain, similar to those seen in the genitalia, are rare.

In rodents, that have stereotyped behavioral sexual dimorphism, several brain regions play canonical roles in sexual behaviors and display sex-specific characteristics. These include the bed nucleus of the stria terminalis (BNST)^9^, anteroventral periventricular nucleus (AVPV)^5,12^, medial preoptic area (MPOA)^5,12^, the ventral medial hypothalamus (VMH)^13^, and the medial amygdala (MeA)^7,14–16^. The MeA, and especially its posterodorsal part (MeApd), was shown to display anatomic and synaptic dimorphism^14,17^ and govern both sexual and pup-directed behaviors^7,15^. MeApd receives direct inputs from both the main and the accessory olfactory bulbs^18^ and expresses receptors for hormones such as estrogen^16^, prolactin^19^, calcitonin^20^ and oxytocin^21^. Therefore, the MeApd is a main candidate to integrate input of social and sexual cues.□

Immediate early genes (IEG) are rapidly transcribed in neurons following increased neuronal activity and are often used to identify populations of active neurons ^22,23^. These genes provide a link between neuronal activity and transcriptional changes. Although IEG are widely used to identify neurons that are active during a specific task, differences in IEG expression can also be observed between neuronal populations even in the absence of an explicit task. Here, we used targeted recombination in active population (TRAP)^24,25^, a genetic reporter for active neuronal populations represented by elevated expression of cFos, to search for sexual dimorphism in baseline activity. We describe a small cluster of cells within the MeApd that is labeled by the TRAP system in females but not in naïve males. This population is labeled in juvenile males before weaning and in adult males after mating or following prolactin treatment. The binary nature of cFos activity in this cluster provides additional insight into the mechanisms regulating sexually dimorphic behaviors.

## Results

We aimed to identify differences in cFos expression in neuronal populations of males and females in brain areas involved in sexual behavior. For this purpose, we used the Trap2 mouse ^24,25^, driving tamoxifen-induced Cre under the cFos promoter (while enabling normal expression of cFos protein) and crossed it with Cre-dependent TdTomato reporter line Ai9 (Figure 1a,b). The Trap2 system allows for permanent recombination, and labeled cells represent high levels of cFos at the time of tamoxifen induction. We first induced recombination and expression of the TdTomato reporter by tamoxifen gavage in young adult males and females (2-3 months old), and mounted brain slices to detect dimorphic populations.

**Figure 1.**
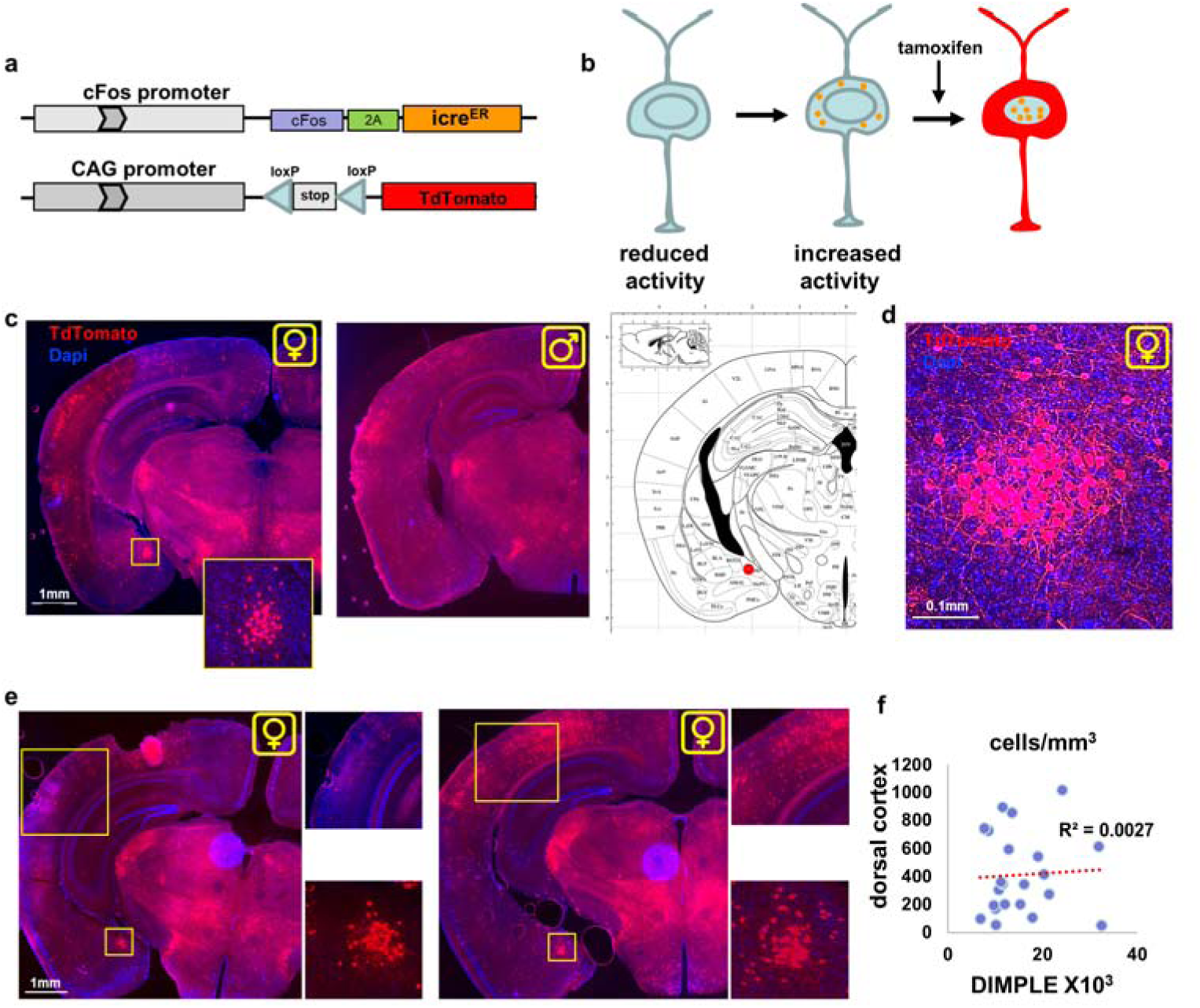
The Trap2 system reveals a sexually-dimorphic neuronal cluster in the medial amygdala. (**a**) Diagram of the Trap2 (cFos-Cre^ER^) and Ai9 (cre-dependant TdTomato) transgenic mice. (**b**) Cre^ER^ is expressed and transported to the cytoplasm upon upregulation of the cFos promoter. Delivery of tamoxifen leads to nuclear translocation of Cre^ER^ and initiates removal of the stop codon and cytoplasmic distribution of TdTomato. (**c**) Screening for sex differences in labeled neuronal population highlighted a cluster of cells (named DIMPLE) in the MeApd that appears in females only. Reference brain image was taken from Paxinos and Franklin’s Brain Atlas ^26^ where the red dot represents the location of DIMPLE. Coordinates: AP- -2.1; ML- 2; DV- 4.8. (**d**) High magnification confocal image of a female DIMPLE. (**e**) Images of females’ brains with low levels (left) or high levels (right) of recombination in other areas of the brain (note dorsal cortex). (**f**) no correlation between the density of cells labeled in the DIMPLE and the dorsal cortex. n=22 images taken from 10 females (2-3 images per animal) R=0.1158 P=0.6080 (spearman). (for higher resolution images see end of document)

### DIMPLE presents high cFos levels exclusively and constantly in the brain of naïve adult females but not males

We found a small (∼200µm in diameter, see measurements in Supplementary Figure 1) cluster of cells located at the MeApd (Figure 1c). This cluster was evident in all females but not in any of the males. We named this cluster DIMPLE according to its bilateral resemblance to facial dimples. Confocal high-resolution images highlight a dense cluster of cells that have local symmetric neurites (Figure 1d). This neuronal cluster was sufficiently distinct that its presence or absence could be readily determined by direct observation under the microscope.

We screened the brains of 96 sexually naïve males and 120 females. Strikingly, all females exhibited DIMPLE labeling via TRAP2, while none of the males did. Importantly, the cohort of 120 females is presumed to include individuals across all stages of the estrous cycle. The absence of a correlation between DIMPLE size—defined by the number of labeled cells within its volume—and the number of trapped neurons in other brain regions suggests that the slight variations in DIMPLE cell numbers observed in females are unlikely to result from technical variability (Figure 1e,f). In males, cells within the MeApd were uniformly distributed, with no apparent clustering.

### Removal of the gonads did not alter the pattern of DIMPLE expression

We next aimed to investigate the role of gonadal hormones in the binary pattern of DIMPLE labeling. We performed ovariectomy in females and orchiectomy in males at two months of age. One-month post-surgery - allowing time for the withdrawal of circulating gonadal hormones -we administered tamoxifen to induce *cFos*-dependent labeling, and animals were sacrificed one month later (Figure 2a). In males, we confirmed the effectiveness of castration by observing degeneration of the seminal vesicles (Figure 2b), consistent with previous reports^27^. However, this hormonal manipulation did not alter the binary pattern of DIMPLE labeling: the cluster remained evident in all females (n=8) and absent in all males (n=5) (Figure 2c).

**Figure 2.**
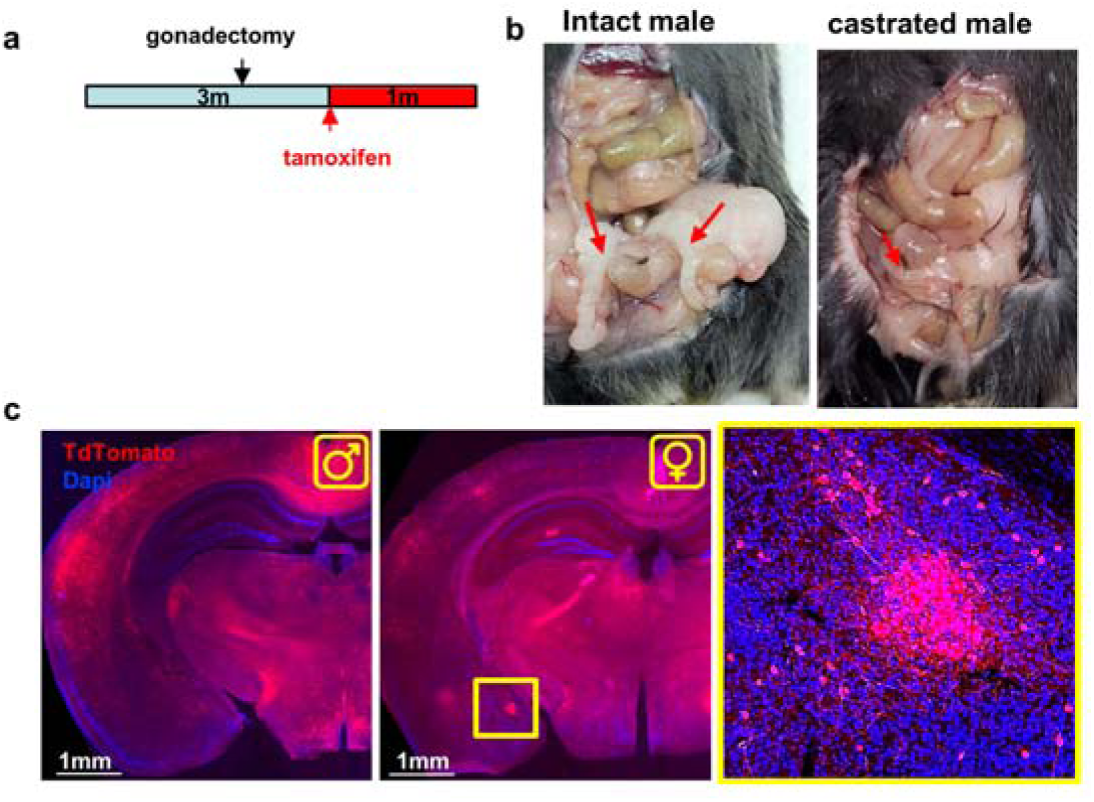
Removal of gonads does not alter the sexual dimorphism of DIMPLE. (**a**) Experimental scheme. Surgery was done at the age of 2 months. Tamoxifen was given at 3 months and brains were retrieved at 4 months. (**b**) The seminal vesicles of castrated males were highly degenerated, verifying the withdrawal of androgens (arrows). (**c**) Examples showing that DIMPLE did not appear in castrated males and was evident in spayed females (enlargement on right).

### DIMPLE is labeled in infant mice of both sexes and disappears in males after weaning

Next, we investigated whether DIMPLE labeling in gained in females or lost in males during postnatal development. To address this, we administered tamoxifen to adolescent pups at postnatal day 24 (p24). One cohort was separated from the mother and any opposite-sex siblings three days afterwards, at P27 (n=15), while the other cohort was separated three days before tamoxifen administration, at p21 (n=9) (Figure 3a). All brains were collected in adulthood at two months of age. We found that DIMPLE labeling in males was observed only in those that remained with the mother and female siblings during the period of tamoxifen exposure (Figure 3b, left). Since all mice received tamoxifen at the same age (p24), these results suggest that DIMPLE activity in males at this stage is contingent on the social environment, likely the presence of the mother or female siblings, rather than developmental age alone. DIMPLE was labeled in all females, irrespective of the timing of weaning (Figure 3c).

**Figure 3.**
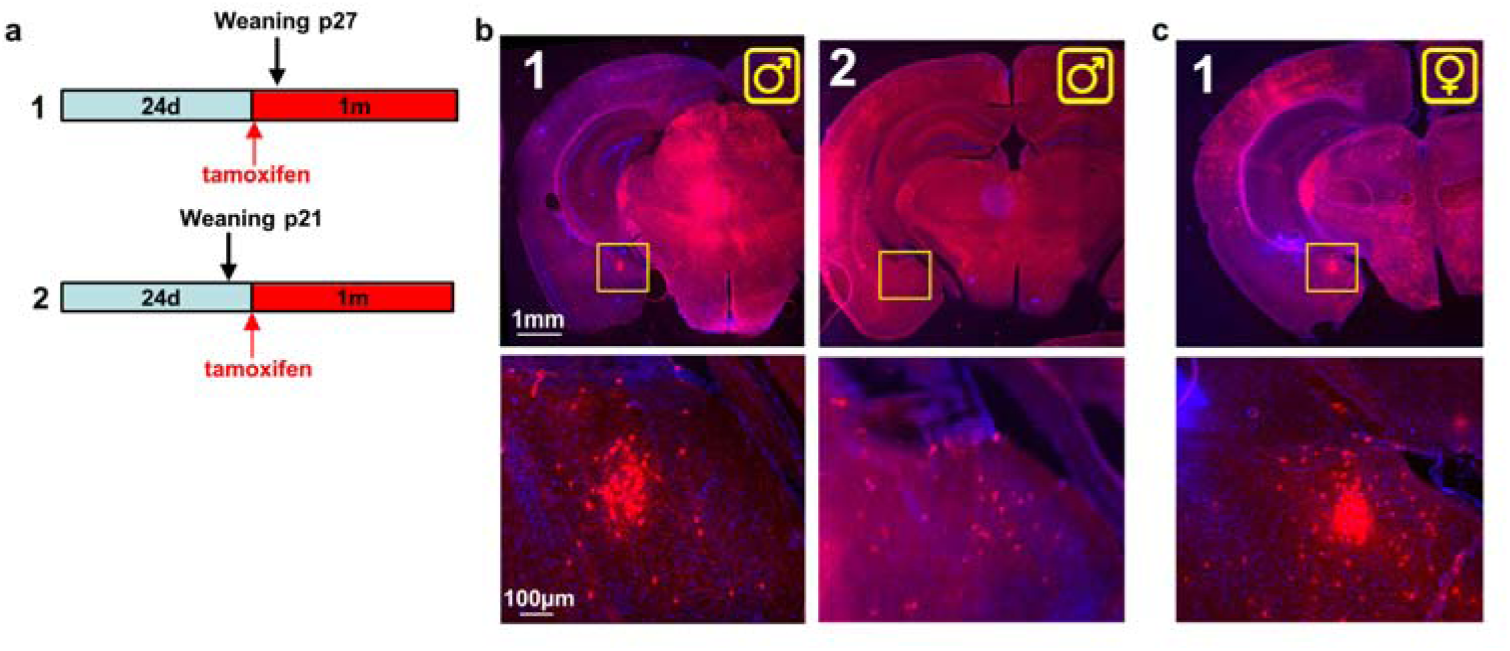
DIMPLE activity during weaning and puberty. (**a**) A scheme of experimental timeline. In both groups, pups were given tamoxifen at the age of 24 days postnatal. Pups in group 1 were separated from the mother 3 days after tamoxifen induction, at the age of 27 days (n=15). Those of group 2 were separated 3 days before tamoxifen induction, at the age of 21 days (n=9). All brains were retrieved 1 month after tamoxifen induction. (**b**) Example images showing DIMPLE in a male labeled with tamoxifen before weaning (group 1) and no DIMPLE in a male labeled after weaning (group 2). (**c**) Example image showing DIMPLE in a female labeled as pup (group 1). DIMPLE appeared in all females regardless of the age of weaning.

The irreversibility of the TRAP system allowed us to label DIMPLE in males before weaning, enabling to visualize it even during adult stages when it is no longer active (see below).

### Tamoxifen alone does not induce DIMPLE activation

Tamoxifen is an estrogen receptor ligand that is commonly used in transgenic mouse models (including the TRAP system) in which Cre recombinase is fused to a modified estrogen receptor, enabling temporal control of its activity. While tamoxifen is designed to have minimal effects on endogenous estrogen receptors, some off-target actions have been reported^28,29^. To rule out the possibility that cFos activity observed in females was artificially induced by tamoxifen’s interaction with estrogen receptor-rich regions such as the MeApd^30^, we compared cFos immunostaining (to highlight current cFos activity) with TRAP-based labeling. For this, tamoxifen was administered to pre-weaning p24 pups (males and females) five weeks prior to brain collection, allowing sufficient time for its clearance from circulation. This approach allowed us to label DIMPLE neurons also in males (Figure 4a). We then prepared histological slices and immunolabeled them for cFos, allowing visualization of both the TRAP and cFos protein signals (Figure 4b). Notably, DIMPLE is more difficult to detect via cFos immunostaining alone due to a higher background signal in the medial amygdala compared to TRAP2-based labeling. We found that approximately 75% of the TRAP-labeled cells in DIMPLE were also cFos-positive in females, compared to only ∼30% in males (Figure 4c). Furthermore, the density of cFos- immunostained cells within the DIMPLE region was significantly higher in females (Figure 4d). These results also provide evidence for the natural silencing of DIMPLE in males following puberty and weaning, reinforcing the idea that DIMPLE activity is transiently present during lactation stages but then suppressed in the adult male brain.

**Figure 4.**
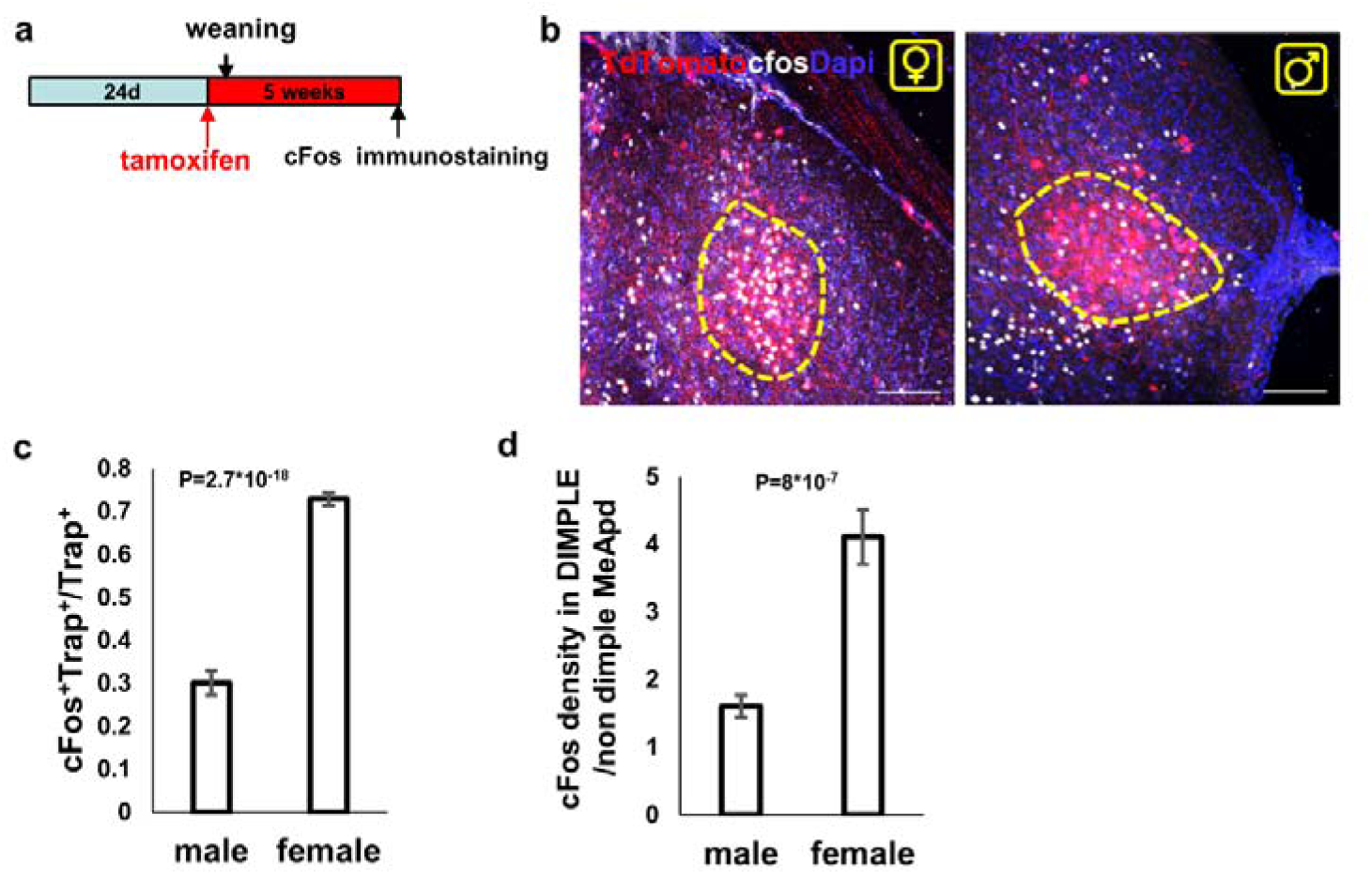
Co-labeling of Trap2 in pre-weaning and cFos in adult. **(a)** A scheme of experimental timeline. (**b**) Images of a male and a female MeApd, DIMPLE is highlighted by dashed line. Scale bar, 100µm. (**c**) Proportion of double-labeled (TRAP□/cFos□) cells relative to the total number of TRAP□ cells within the DIMPLE region. (**d**) Ratio of cFos□ cell density within the DIMPLE compared to the surrounding tissue. Student’s T-test. n=6 males and n=6 females, 4-8 images per animal.

### Sexual activity induces cFos activation in the DIMPLE of males

Given the established role of the MeA in regulating sexual behavior^31^, we next investigated whether DIMPLE becomes active in males following sexual experience. To this end, we co-housed male and female mice and administered tamoxifen to mating pairs upon detection of a vaginal plug. Remarkably, DIMPLE labeling was observed in males under these conditions (Figure 5a) and was, on average, even more prominent than that observed in females or naïve males labeled before weaning (Figure 5b, Supplementary Figure 1). Furthermore, DIMPLE was labeled in a male that had been co-housed with a female for only one hour, during which successful mating occurred, followed by immediate administration of 4-hydroxytamoxifen (the fast-acting tamoxifen metabolite). This resulted in detectable activation (Figure 5c, horizontal view), indicating that DIMPLE responds rapidly following mating.

**Figure 5.**
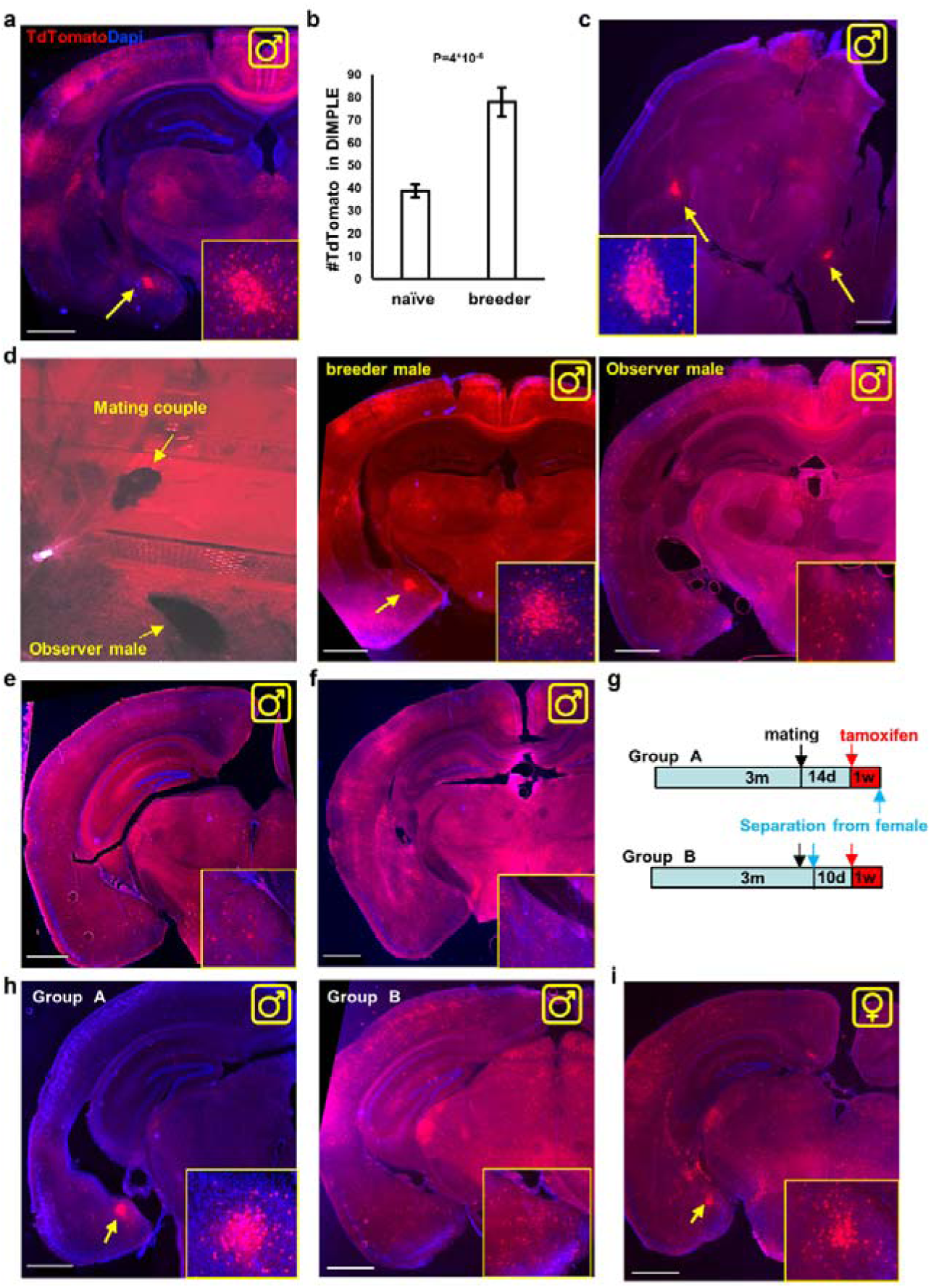
DIMPLE activity during sexual behavior. (**a**) Three-months old males were housed with females. Tamoxifen was given to males at the day of vaginal plug detection. Example image shows evident DIMPLE in males one day after sexual activity. n=11 animals. (**b**) Comparison of the number of TdTomato+ cells in the DIMPLE per 50µm slice in adult naïve males labeled pre-weaning vs. males labeled after mating. n=9 naïve, 6 breeders, 2-6 images/animal. Student’s t-test. (**c**) Horizontal brain slice of a male that managed to mate within an hour after being put with a female. The male was given 4- hydroxytamoxifen immediately, and the brain was retrieved after 1 week. (**d**) Females and males were housed in a cage divided by a barrier, allowing interaction without permitting sexual contact (n=3 males, 2 females). Another male was given full access for mating. Tamoxifen was given to all males 1 day after the female introduction. The mating male had clear DIMPLE labeling while no labeling of DIMPLE was evident in any of the 3 observer males (example images shown) (**e**) Males (n=3) were grown for 24h in a cage with female bedding before tamoxifen administration. Example image shows no labeling of DIMPLE. (**f**) Example image from the brain of a male that was kept with a female but failed to induce pregnancy for >2 weeks. No DOMPLE was observed (n=3). (**g-i**) (**g**) Timeline of experiments testing the stability of sexually-activated DIMPLE. Males were mated with females and tamoxifen was administered to the males 2 weeks after vaginal plug detection either in the continuous presence of females (group A, n=4) or following 10 days of separation from their female (group B, n=4). (**h**) Example images show labeling of DIMPLE in males that were kept together with their female (group A) but not in those that were separated from the female (group B). (**i**) DIMPLE was labeled in all pregnant females from the experiment (11 animals). Note that using tamoxifen in pregnant females can prevent successful delivery and lead to intrauterine death. Scale bar for all images, 1mm.

To determine whether direct sexual contact, rather than the mere presence of receptive female, was necessary for DIMPLE activation, we housed three males and 2 females in a cage separated by a barrier permitting visual, auditory, and olfactory interaction but preventing physical contact. Another male was housed in the females’ section (Figure 5d) to allow mating. While it was strongly labeled in the mating male, in the observer males, DIMPLE was not labeled, (Figure 5d). DIMPLE was also not labeled in males exposed to female-soiled bedding for several days (Figure 5e, n=3), or in males that remained co- housed with females but failed to induce pregnancy (Figure 5f, n=3).

Mouse pregnancy lasts approximately 19 days. To determine how long DIMPLE remains active in males following mating, we administered tamoxifen two weeks after vaginal plug detection. One group of males remained with their female partner throughout this period (Group A, n=4), while a second group was separated from the female four days after plug detection (Group B, n=4) (Figure 5g). DIMPLE labeling was observed in all Group A males -those that remained with the female throughout the pregnancy period (Figure 5h, left) but in none of group B (Figure 5h, right). This suggests that sustained cohabitation with the pregnant female is required to maintain DIMPLE activity in males. As expected, DIMPLE was evident in all females in these experiments, regardless of pregnancy status (Figure 5i).

### Prolactin is sufficient but not necessary for DIMPLE induction

The findings above suggest that DIMPLE is likely to be responsive to a hormone released during mating. To identify candidate hormonal pathways, and for better understanding the gene expression in DIMPLE, we used the Amygdala single-cell RNA sequencing database of Hochgerner et al. ^32^. This dataset includes neuronal clusters of the whole amygdala aligned with anatomical regions through cross-referencing with the Allen Brain Atlas. Using their supplemental online engine, we searched for neuronal populations whose spatial distribution corresponds to the DIMPLE location in the MeApd. We identified several candidate clusters - primarily GABAergic clusters 29–34 that closely match the anatomical site of DIMPLE (Figure 6a). Several receptors for potential hormonal mediators were enriched, including estrogen receptors 1 and 2 (*Esr1* and *Esr2*), the calcitonin receptor (*Calcr*), and most prominently, the prolactin receptor (*Prlr)* (Figure 6b). *Prlr* was also shown by others to be localized to the MeApd^33–35^ (Figure 6c). Given that prolactin is acutely secreted in male rodents following mating^19,36^, these findings position prolactin as a strong candidate for mediating DIMPLE activation in males. Since the data of ^32^ and the Allen brain atlas in-situ are both from males, we performed bulk RNAseq of MeApd slice containing the DIMPLE in both virgin males (induced at p24 before weaning to visualize the DIMPLE) and females (n=3 females, 4 males, supplementary table 1 and GSE297014). We found no significant changes in *Prlr* reads between males and females (Figure 6d). *Prlr* has not been listed as a sexually-dimorphic gene in the MEA in other studies^7,37^, leading to the hypothesis that prolactin itself, and not the receptor levels, is dimorphic, as supported also by literature^38^ . We therefore tested whether prolactin is sufficient to induce DIMPLE in naïve males. We administered intraperitoneal prolactin to naïve adult males (n=4), followed by tamoxifen induction one hour later. DIMPLE labeling was observed in these animals (Figure 6e and supplementary Figure 1), indicating that systemic prolactin alone is sufficient to activate this region in males in the TRAP system. These results support the hypothesis that prolactin released during mating may be the physiological trigger for DIMPLE activation in males.

**Figure 6.**
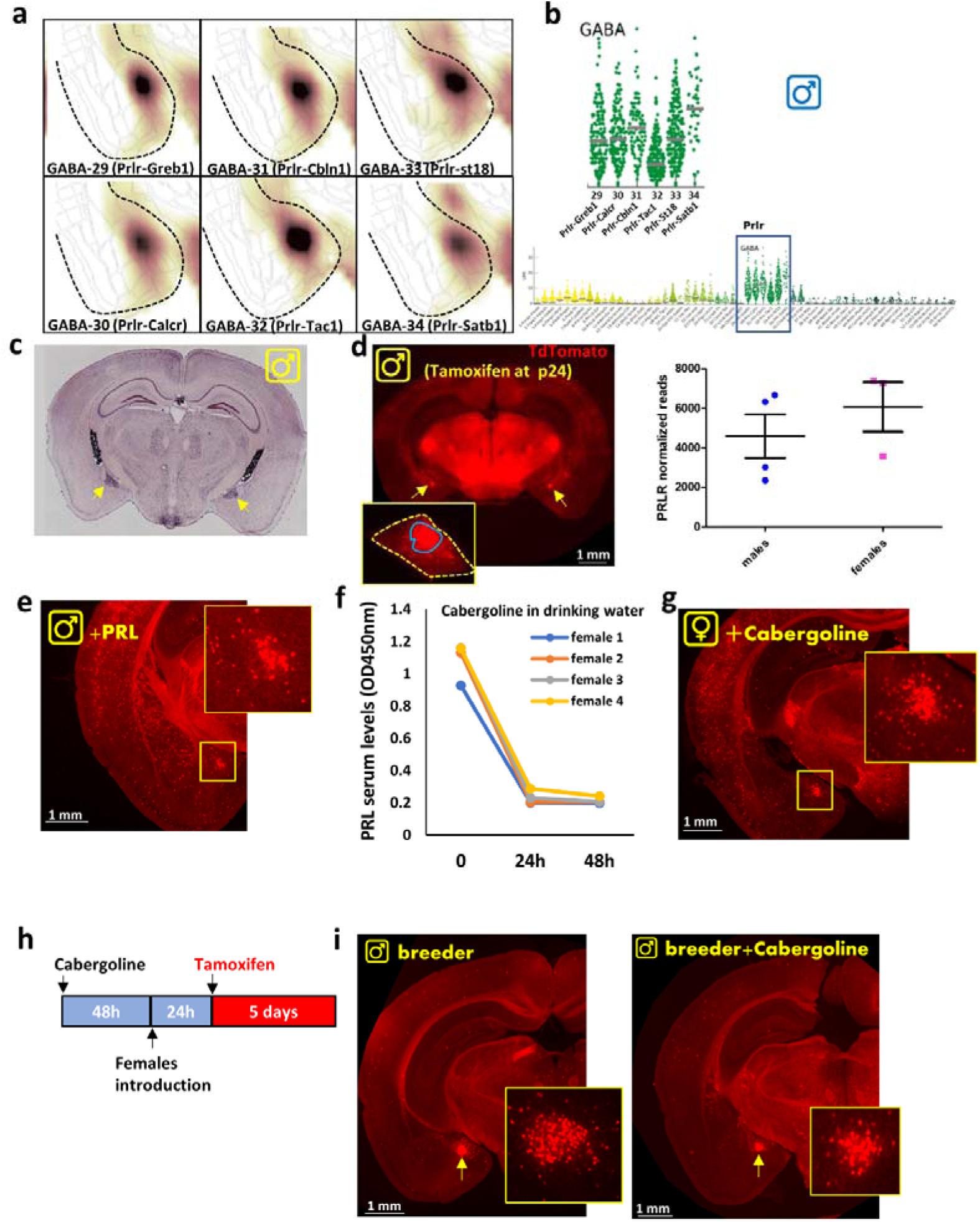
Prolactin signaling and DIMPLE activation. (**a-b**) Single-cell RNA sequencing of the male amygdala from^32^ reveals (**a**) GABAergic clusters that map to the MeApd in a region resembling the location of DIMPLE. Dashed line highlights the amygdala-hypothalamus border. (**b**) The two-gene identifier and the violin plot for *Prlr* demonstrate specific enrichment of this gene within those clusters. Each dot represents *Prlr* levels in one cell. (**c**) In-situ hybridization for *Prlr* in the MeA (arrows) taken from the Allen Brain atlas (exp. 72340223). (**d**) Bulk RNA-seq on manually dissected MeA tissue (outlined by the dashed yellow line), which included the DIMPLE region (indicated by the blue line) in males and females (n=4 and 3). Approximately 20% to 40% of the dissected area encompassed the DIMPLE. We labeled males before weaning at p24 and extracted the tissue in the adult. In DESeq2 analysis, *Prlr* levels were insignificant between males and females. For significant differentially-expressed genes, see supplementary table 1. (**e**) An example of DIMPLE labeling in a male that was treated with Prolactin IP and given tamoxifen 1h later. DIMPLE was evident in all mice (n=4). (**f-g**) Females (n=8) were treated with cabergoline to inhibit pituitary prolactin release and labeled with tamoxifen 24 and 48h later. (**f**) ELISA analysis of blood samples from 4 females confirmed effective suppression of prolactin following cabergoline treatment. (**g**) An example of DIMPLE labeling in cabergoline-treated females. DIMPLE labeling was evident in all 8 females with this treatment. (**h-i**) The effect of cabergoline on DIMPLE labeling in breeder males. (**h**) Males (n=3) were given cabergoline in their drinking water, females were introduced 2d later (each pair housed together in a single cage). Tamoxifen was administered 24h later, and the males and females remained paired in the same cage, continuing to receive cabergoline until brain retrieval, 5 days later. Three additional males underwent the same protocol, but without cabergoline treatment. (**i**) Example images of DIMPLE (arrows) in breeder males with and without cabergoline.

To determine whether prolactin is necessary for DIMPLE activation in females, we inhibited endogenous prolactin secretion using cabergoline, a potent dopamine receptor agonist that suppresses prolactin release via action on hypothalamic dopaminergic neurons. Cabergoline was administered continuously in the drinking water. Successful suppression of circulating prolactin levels was confirmed via ELISA analysis (Figure 6f). Tamoxifen was administered either 24 or 48 hours after initiation of cabergoline treatment. Despite the significant reduction in prolactin, DIMPLE labeling remained evident in all treated females (n=8), suggesting that prolactin is not required for DIMPLE activation in females under basal conditions (Figure 6g). We tested whether prolactin inhibition could suppress DIMPLE activation in breeder males. Males (n=3) were given cabergoline in their drinking water, and two days later, females were introduced into the males’ cages, with each pair housed together in a single cage. Tamoxifen was administered 24 hours later, and the males and females remained paired in the same cage, continuing to receive cabergoline until brain retrieval, 5 days later (Figure 6h). Three males underwent the same protocol but without cabergoline treatment. Cabergoline was ineffective in inhibiting DIMPLE activation in breeder males, as it was observed in the brains of all animals in this experiment (Figure 6i). We conclude that prolactin is involved in only part of a more complex set of processes that drive brain changes in males following mating. Our RNA-seq data (supplementary table1 and GSE297014), along with previous studies^7,37^ highlight additional sexually dimorphic genes in the MeApd, some of which consistently appeared as sexually dimorphic across all analyses: *Brs3, Ednra, Fgf11, Foxq1, Fxyd6, Igfbpl1, Kcng2, Nfix, Npbwr1, Pdlim1, Pgr15l, Prokr2, Rbm20,* and *Rrad*. These demand further investigation, together with hormonal profiling for additional mediators.

## Discussion

Sexual dimorphism in the brain is an area of increasing research interest. In rodents, whose sexual behavior is highly stereotyped^11,39^, sex differences have been observed across brain anatomy, circuitry, gene expression, and hormonal responsiveness. In this study, we examined sex differences in the baseline expression of the immediate-early gene cFos in neurons. We report that a distinct cluster in the MeApd, which we termed DIMPLE, displays robust and reproducible binary dimorphism. DIMPLE was labeled in all females but not in adult virgin males, showing 100% reproducibility across sexes. Furthermore, DIMPLE activity serves as a stable marker of the social or reproductive state in males, as it is active during lactation, de-activates after weaning, and reactivates following sexual experience. Finally, we demonstrate that prolactin is sufficient to induce DIMPLE labeling; however, blocking prolactin signaling was not sufficient to suppress or reverse this activation.

Many studies have examined the neural changes that occur in preparation for parenthood, more commonly in females ^33,40–42^, but also in males^19,43–46^. It is well established that male rodents exhibit a behavioral shift following mating, from aggression toward pups to parental behaviors such as grooming, pup retrieval, and nest incubation ^43,45,47,48^. and part of this change has been attributed to GABAergic neurons in the MeA^7^.

The finding that cFos is activated in the male DIMPLE shortly after sexual contact, and remains active for as long as the male is with the female, supports the hypothesis that DIMPLE may be part of the neural circuitry that suppresses infanticidal behavior. Notably, DIMPLE activity disappears if the male is separated from the female, highlighting its dynamic and state-dependent nature. Similarly, adolescent males may suppress infanticide behavior when co-housed with their mother and younger siblings. Previous studies have implicated the bed nucleus of the stria terminalis (BNST)^45,49^ , the medial preoptic area (MPOA)^50^ the amygdalo-hippocampal area^51^ and ventromedial hypothalamus (VMH)^13^ in mediating the shift from infanticidal to parental behavior. The MeA, particularly the MeApd also plays a role in this circuitry^7^. Whether DIMPLE acts as a marker of this functional state or actively contributes to it remains to be determined. The MeApd projects robustly to both the BNST, MPOA and VMH^52^, suggesting that DIMPLE neurons may lie upstream of these established regulators of parental behavior. Nevertheless, DIMPLE is activated immediately after mating, much earlier than other reports of parental-related changes^48,53^. Therefore, the activation of DIMPLE may also be associated with other, yet-to-be-identified processes distinct from the established timeline of paternal adaptations. Targeting DIMPLE to investigate its functional roles is an important yet challenging direction for future research, given its small size and the lack of a unique molecular handle.

Given that DIMPLE is activated in males following mating, hormones released during sexual activity likely contribute to its activation. Among potential candidates, we highlight prolactin as a potential mediator. The prolactin receptor (PRLR) is expressed in the MeApd, as shown by single-cell RNA-sequencing data^32^, transgenic lines^19,34^, and in-situ mapping from the Allen Brain Atlas^35^. In line with this, our findings demonstrate that systemic administration of prolactin in virgin males is sufficient to induce DIMPLE activity. Smiley et al. ^19^ reported activation of PRLR-expressing neurons in the MeApd and other brain regions following exposure of virgin males or sires to pups. Furthermore, conditional knockout of PRLR in Vgat- or CKC-expressing neurons led to deficits in paternal behaviors, underscoring the functional relevance of prolactin signaling ^19^. Notably, Smiley et al. also showed that prolactin levels peak in male serum immediately after ejaculation but decline rapidly thereafter. This transient prolactin surge is not essential for the expression of paternal behaviors. Instead, sustained circulating prolactin levels in males cohabiting with the dam and pups are the ones required for ongoing paternal care^19^. We demonstrate that DIMPLE remains active as long as the male stays with the female, but is silenced upon separation. However, our findings regarding prolactin involvement are somewhat ambiguous; while prolactin alone was sufficient to induce DIMPLE activation in virgin males, post-mating males with prolactin inhibition still showed DIMPLE activation. We do not rule out the involvement of additional factors, particularly in females, where suppression of prolactin did not abolish DIMPLE activation as well. For example, calcitonin receptor is enriched in the MeApd in females and has a role in social behavior ^20^.

In conclusion, the all-or-none pattern of DIMPLE activation - present in all females but only in parental and pre-weaning males - suggests that it may function as a switch within a neural circuit governing sexual maturation and post-mating behavior. Findings such as these contribute to our understanding of the neural regulation of complex social behaviors and the mechanisms underlying sexual dimorphism in the brain.

## Methods

### Mice

All animal procedures were approved by the animal care and use committee of the Hebrew University. Both males and females were used. Age and sex of animals are indicated in the relevant section of the results. Transgenic mouse lines used in this study were as follows: Trap2^24,25^ and Ai9 ^54^ were purchased from Jackson Laboratories (RRID: IMSR_JAX:030323 and IMSR_JAX:007909); these mice are of C57/BL6 background. Tamoxifen (Sigma T5648, 30 mg/ml in sunflower seed oil) was administered by oral gavage at 200 mg/kg. Pups were given tamoxifen orally using a flexible gel-loading pipette tip. 4-hydroxytamoxifen (Sigma H6278) was diluted in saline containing 1% tween-80 and 5% DMSO (Sigma) and was given i.p at 6 mg/kg (Figure 5c only). Mouse Prolactin (Cyt- 321, ProSpec, Israel) was dissolved in saline and given i.p at 120µg/kg. Cabergoline (0.5mg, Trima, Israel) was dissolved in a drinking bottle of 0.5 L. Regular specific pathogen–free housing conditions with a 12-hour light/ 12-hour dark cycle were used. Irradiated rodent food and water were given ad libitum. A Plexiglass cage, sized 50X50cm, with metal separation bars, was used for the experiment described in Figure 5e. For mating experiments, one female and one male were held in every cage.

### Orchiectomy

Males aged 2 months were anaesthetized (ketamine 50 mg/kg and medetomidine 10mg/kg i.p) and analgesia was given (Meloxicam 5mg/kg s.q). Two incisions were made on both sides of the scrotum, and the testicles were pushed out. Blood vessels and the vas deferens were crushed by a hemostat, and the testicle was cut by a scalpel. A suture was made to keep the spermatic cord stump in the abdomen. Mice were woken up by atipamezole (2 mg/kg i.p).

### ovariectomy

Females aged 2 months were anesthetized using isoflurane, and analgesia was administered by meloxicam (5 mg/kg, s.q.). The back was shaved, and a 2 cm skin incision posterior to the rib cage was made. Blunt dissection through the abdominal muscles was performed to access the periovarian fat pad on each side. The ovary was carefully exteriorized, and the associated blood vessels and fallopian tube were crushed using a hemostat. The ovary was then excised with a scalpel. The tissue was returned to the abdominal cavity after confirming the absence of active bleeding. Finally, the muscle and skin layers were closed using tissue clips.

### RNAseq

Tamoxifen was given to 3 females at 6w and to 4 males at p24 before weaning. At 8w, Animals were anesthetized with 5% isoflurane and decapitated, and 300 µm coronal slices were made by vibratome (Leica) in ice-cold aCSF solution equilibrated with 95% O_2_/5% CO_2_ ^55^. Slices containing the MeA were inspected under a fluorescent stereoscope (Nikon SMZ25) and the DIMPLE area of both hemispheres was carefully dissected, imaged to evaluate % DIMPLE in the dissected tissue, and immediately processed by Zymo Quick- RNA™ Microprep kit (cat#R1050). mRNA library preparation and sequencing were carried out by the Genomic Applications Laboratory at the Core Research Facility of the Hebrew University, following a standard protocol. RNA quality was assessed using the TapeStation 4200 system (Agilent Technologies), High Sensitivity RNA ScreenTape (5067-5579) along with the Qubit® RNA HS Assay Kit (Invitrogen; Q32852). For library preparation, 10 ng of RNA from each sample was processed using the SMARTer® Stranded Total RNA-Seq Kit v3 - Pico Input Mammalian (Takara). Samples were eluted in 25μL of 10 mM elution buffer, then all of them were pooled into 10nM final concentration into a single tube. The library was sequenced using NovaSeq6000 machine (Illumina) with 100 cycles and single-Read sequencing conditions. The NextSeq basecalls files were converted to fastq files using the bcl2fastq program with default parameters. Raw reads were quality-trimmed with cutadapt using a quality threshold of 32 for both ends, poly-G sequences (NextSeq’s no signal) and adapter sequences were removed from the 3’ end, and poly-T stretches were removed from the 5’ end (being the reverse-complement of poly-A tails). The cutadapt parameters included using a minimal overlap of 1, allowing for read wildcards, and filtering out reads that became shorter than 15 nt. Finally, low quality reads were filtered out using fastq_quality_filter, with a quality threshold of 20 at 90 percent or more of the read’s positions. Processed reads were aligned to the mouse GRCm39 genome with Tophat, v2.2.1, allowing for 5 mismatches, using gene annotations from Ensembl release 106. Raw counts per gene per sample were obtained with htseq-count, v2.0.1, then normalization was done with the DESeq2 package. The significance threshold was taken as padj<0.1 (default). Raw and processed data (DeSeq2) of RNA sequencing can be found at GSE297014.

### Prolactin ELISA

For prolactin measurement, animals were gently warmed under red light to facilitate vasodilation. Blood samples were collected from the tail vein into 1□mL EDTA-coated tubes. Samples were centrifuged at 1,000□×□g for 10 minutes at 4□°C, and the plasma was separated and stored at –20□°C until all experimental time points were collected. Plasma prolactin levels were measured using a mouse prolactin ELISA kit (Wuhan USCN SEA846Mu). For each sample, 20μL of plasma was diluted to a final volume of 100μL using the kit’s dilution buffer, and the assay was carried out according to the manufacturer’s instructions. Samples were read at 450 nm with a reference value of 540 nm using a Tecan Infinite f200 Pro 96-well plate reader.

*Histology and Immunofluorescence*.

Brains were fixed by immersion in 4% paraformaldehyde on ice for 12h, washed with PBS, and sectioned into 50-μm coronal sections by a vibratome (Leica). Immunostaining was done as described ^56^ with anti-cFos (1:300; synaptic systems RRID: AB_2231974). Alexa 647 anti-rabbit IgG (RRID: AB_2492288) was from Jackson Immunoresearch (1:400 dilution). Sections were mounted on glass slides and covered with mounting medium containing DAPI (SouthernBiotech).

### Microscopy

Confocal microscopy was done using Olympus FV-1000 on 10X, 20X and 40X objectives and 2 µm distance between confocal z-slices. Low-magnification images were acquired using a Nikon SMZ-25 fluorescent stereoscope on X1 and X2 objectives.

### Measurements

DIMPLE diameter and cell density were measured using *ImageJ* and *Olympus FV10-ASW Viewer*. To calculate cell density, the area of the DIMPLE or a reference region (dorsal cortex; ∼1.5 × 1□mm encompassing all six cortical layers) was measured and multiplied by the slice thickness (50□µm) to determine the volume. Cell counts within these regions of interest (ROIs) were performed manually by an experimenter blinded to experimental conditions. Double-labeling of TRAP and cFos was assessed manually using the 3D projection tool of the *Olympus FV10* viewer. cFos expression in the DIMPLE and adjacent medial amygdala was quantified using a custom-written MATLAB script. Regions of interest (ROIs) were manually delineated, with care taken to exclude areas lacking tissue. The statistical analysis method used is described in the corresponding experimental section.

## Acknowledgements

We thank Drs. Abed Nasereddin, Idit Shiff, Yuval Nevo and Inbar Plackes from the Genomic Applications Laboratory, Core Research Facility at the Hebrew University for their support with RNA sequencing. Prof. Adi Mizrahi and Dr. Baruch Haimson for mice and helpful advice.

---

**Supplementary Figure 1:**
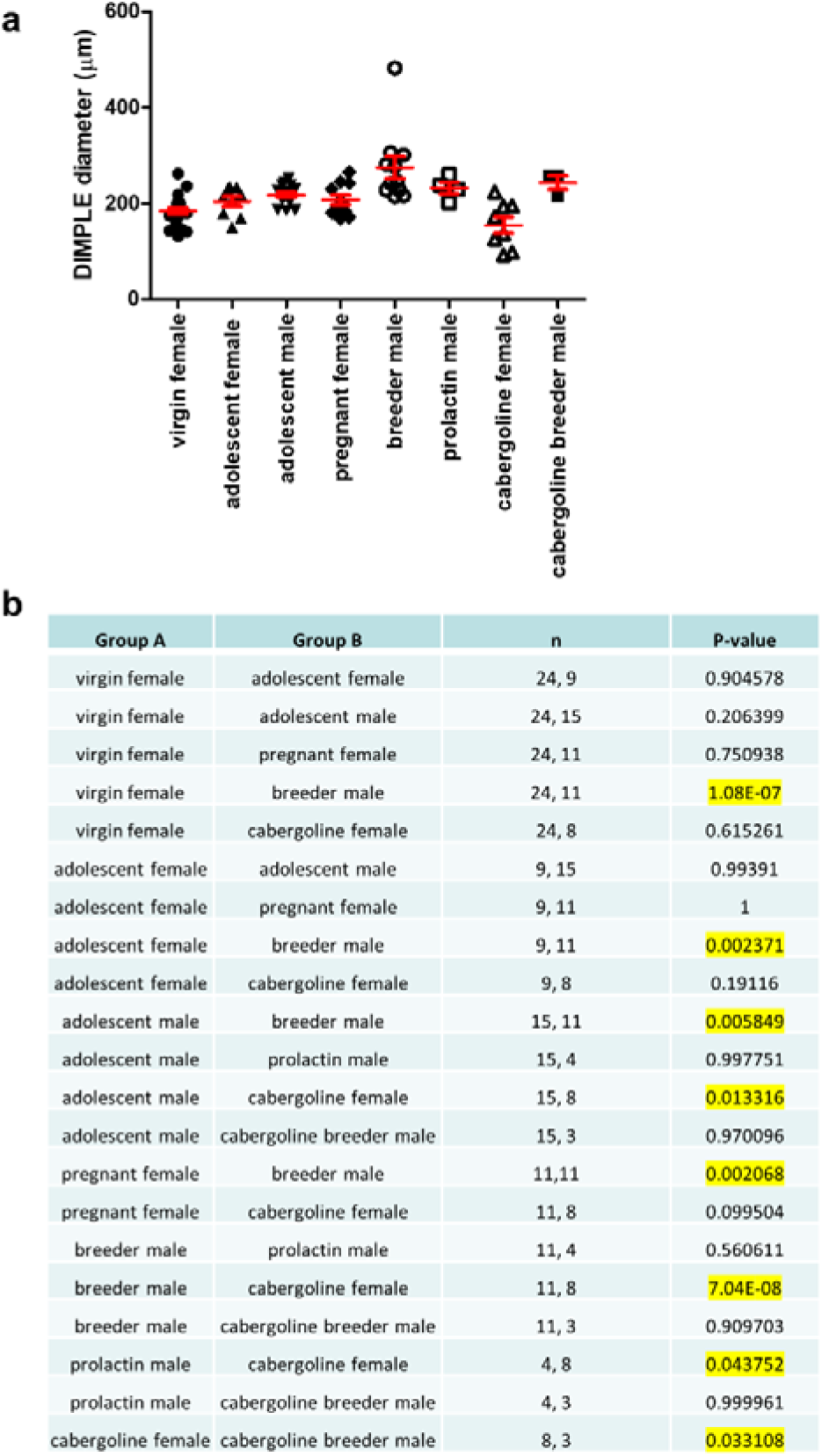
(**a**) Measurements of DIMPLE diameter (in µm) and (**b**) results of a one-way ANOVA test showing the significance of differences between all groups. **Notes**: 1. The virgin male group is not represented in the graph as no cluster of TdTomato cells was detected in these animals (n=96), making the difference highly significant compared to all other groups. 2. Adolescent males and females refer to adult animals that received tamoxifen during late lactation. 3. Group p-value = 3.1 × 10□□.

