## Supplementary figures and images for "A Sexually Dimorphic Neuronal Cluster in the Mouse Medial Amygdala Exhibits Binary Activation Mode Based on Male Sexual Status"

### SUPPLEMENTARY FIG1

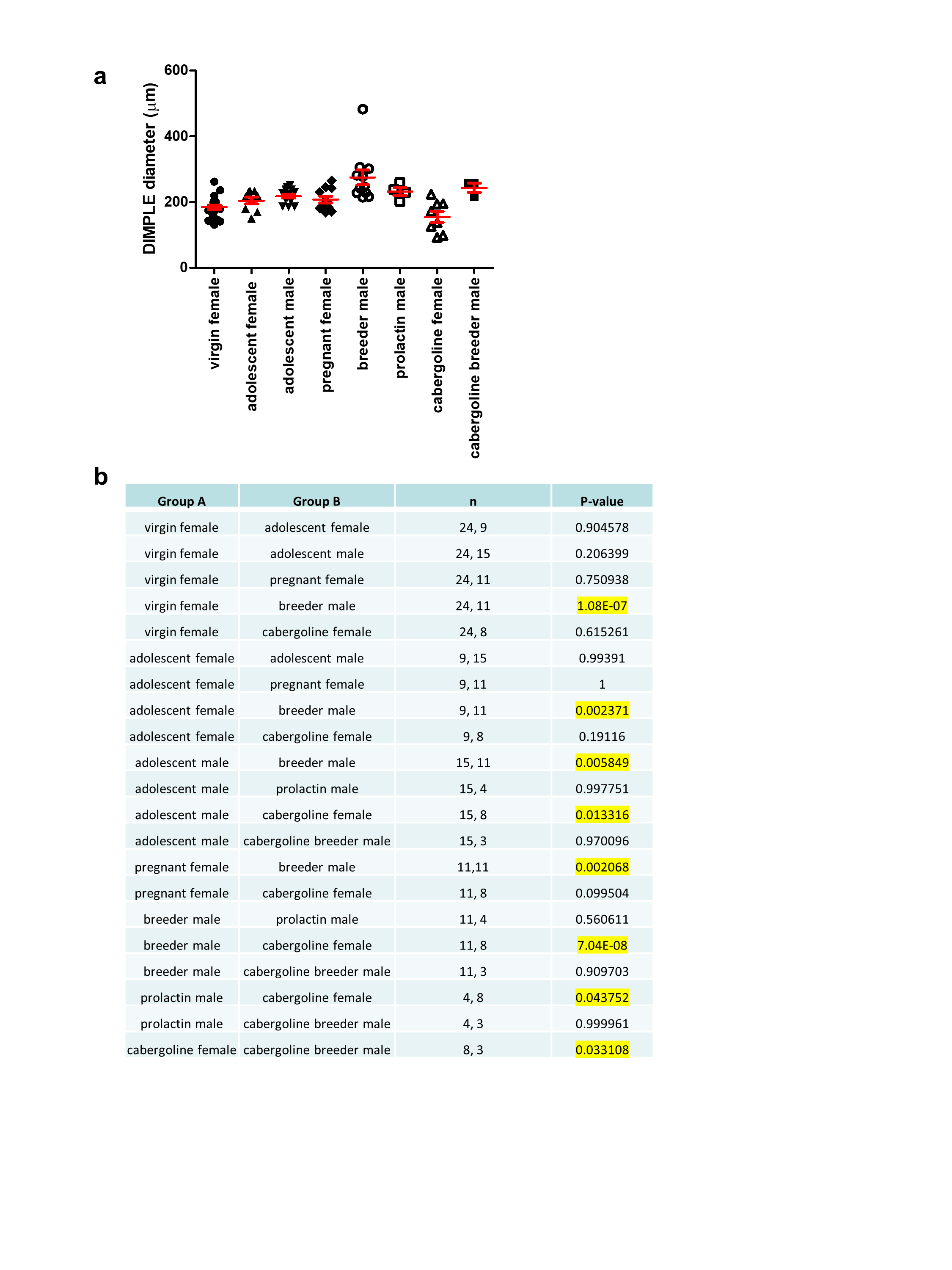
